# Mechanistic insights of radiation-induced endothelial senescence impelling glioblastoma genomic instability at relapse

**DOI:** 10.1101/2021.12.13.472364

**Authors:** Charlotte Degorre, Ophélie Renoult, Ann Christin Parplys, Hala Awada, Anne Clavreul, Manon Pietri, Lisa Oliver, Noemie Joalland, Michelle Ricoul, Catherine Gratas, François Vallette, Kirsten Borgmann, Laure Sabatier, Claire Pecqueur, François Paris

## Abstract

Despite aggressive clinical protocol, all glioblastoma (GBM) recur at the initial site within the irradiated peritumoral microenvironment. Whereas irradiated microenvironment has been recently proposed to accelerate GBM relapse, molecular and cellular mechanisms remain unknown. Here, using relevant *in vitro* and *in vivo* models, we decipher how radiation-induced endothelial senescence drives the emergence of aggressive GBM cells. Secretome (SASP) of radiation-induced senescent (RIS) endothelium enhances genomic instability and intratumoral heterogeneity in irradiated GBM cells. In-depth molecular studies revealed that CXCL5 and CXCL8, from the SASP, activate CXCR2 receptor on tumor cells leading to increased DNA hyper-replication, micronuclei formation and aneuploidy. Importantly, through CXCL5/8-CXCR2 axis activation, this SASP increases GBM aggressiveness *in vivo*. Both chemokines were detected in relapsing, but not primary, GBM biopsies and positively correlated with worst patient outcome. In conclusion, we identify new molecular and preclinical insights of relapsing GBM aggressiveness where RIS vascular niches fuel aggressive tumor emergence.

## Introduction

Radiotherapy (RT) in combination with surgery and chemotherapy remains the gold standard treatment for patients with glioblastoma (GBM) ^1,2^. RT efficacy is challenged by intrinsic tumor resistance of infiltrating GBM cells imposing large fields of RT with 2-3 cm margins around the tumor leading to significant exposure of the surrounding healthy tissue ^3^. Despite those aggressive treatments, overall median survival of GBM patients does not exceed 18 months, with all GBM recurring within a year. Management of recurrent GBM, currently limited to palliative care, urgently needs new perspectives of treatment that will come from a better understanding of molecular mechanisms involved in GBM relapse ^4^. Surprisingly, 90% of GBM recurrences occur at the surgery margins, wherein the peritumoral brain tissue has been irradiated. This singularity has been explained by the acquisition of radiation resistance and growth properties of surviving GBM cells ^5^. Radiation usually triggers tumor cell death following the massive formation of DNA damages. When tumor cells survive, mis-or un-repaired DNA damage leads to nucleotide alterations and chromosomal rearrangements, contributing to whole-genome remodeling. By conferring diverse fitness advantages to tumor cells, genomic remodeling gives rise to multiple and heterogeneous subpopulations of RT-surviving GBM cells and contributes to intratumoral heterogeneity, which has been described as the root cause of tumor relapse aggressiveness and resistance to treatments. A recent study proposed a different hypothesis where pre-irradiation of the brain accelerates GBM relapse in orthotopic mouse models, in particular through radio-induced astrocyte senescence ^6^.

Senescence is a premature aging process characterized by cell cycle arrest that promotes tissue remodeling during embryonic development. In the case of oncogenic damages, senescence limits tumor progression ^7^. However, at the same time, senescence can also promote inflammation and cancer development in aged organisms ^8,9^. While senescence programs depend on cell type and stress origin, all senescent cells display singular phenotypic features including a singular secretome known as senescence-associated secretory phenotype (SASP) ^10^. This SASP is characterized by pro- and anti-inflammatory signaling molecules, proteolytic enzymes and components of the extracellular matrix ^8^. While astrocytes are prone to radiation-induced senescence, vascular remodeling has been described as a hallmark of ionizing radiation injury, in particular through endothelial cell apoptosis or senescence ^11^. Importantly, in post-mortem biopsies of recurrent GBM in patients following 60Gy, senescence has been specifically detected in brain endothelial cells ^12,13^. Thus, the mechanistic understanding of how radiation-induced endothelial senescence may affect tumor behavior is crucial to limit GBM recurrence.

In this study, we investigate the involvement of senescent endothelial cells in RT failure and GBM relapse. Using our previously established Radio-Induced Senescent (RIS) model of quiescent human primary microvascular endothelial cells ^14^, we demonstrate in the present study that the SASP of RIS endothelial cells enhances radiation-induced genomic instability and intratumoral heterogeneity in GBM cells, resulting in very aggressive GBM progression *in vivo*. Using relevant and diverse *in vitro* and *in vivo* models, we demonstrate that SASP alters DNA replication, and exacerbates abnormal division, micronuclei formation and aneuploidy triggered by irradiation. We further identified CXCL8 and CXCL5 in SASP of RIS endothelial cells, and their shared receptor CXCR2 as key players mediating SASP-induced genomic instability, using specific pharmacological inhibition of each actor, biochemical analyses and murine orthotopic models of GBM. Finally, we provide clinical relevance of our findings since both cytokines were detected in recurrent GBM biopsies after Stupp protocol. Our results prove that radiation creates senescent niches that directly impel GBM relapse and aggressiveness.

## Results

### Secretome of RIS endothelial cells drives the emergence of aggressive radiation-surviving GBM cells

We exploited our recently published human primary endothelial model of RIS to determine the impact of endothelial cell secretome on parental human U251 GBM cell response to radiation and characterize corresponding radiation-surviving GBM cells (*Figure 1a & Table 1*). Briefly, clonogenic assays were performed using U251 GBM cell line preconditioned for 18 hours with the secretome of either unirradiated (CM) or RIS endothelial HMVEC-L cells (SASP) in response to escalating doses of irradiation. The SASP did not affect U251 cell plating efficiency (*Supplementary Figure 1)* nor radiation sensitivity as shown by the similar number and size of colonies counted after radiation (*Figure 1b-c*). Then 15Gy-surviving U251 cells cultured in either CM or SASP (designed hereafter as R15CM and R15SASP respectively) were collected and further analyzed using both phenotypic and molecular analyses. Surprisingly, and despite similar proliferation rate *in vitro* (*Figure 1d*), orthotopic injection of R15SASP cells into the cerebral subventricular zone of NSG mice triggered the development of more aggressive tumors than R15CM cells as shown by reduced tumor-bearing mice survival (*Figure 1e*, median survival of R15CM: 58 days *vs* R15SASP: 42 days, p=0.0028). RNA sequencing analysis using DGEseq technology clearly revealed a distinct molecular pattern between R15CM and R15SASP cells (*Figure 1f*). However, further bioinformatic investigation did not identified specific biological networks (*Data not shown*). To investigate the molecular heterogeneity of radiation-surviving U251 cells, individual clones were collected from clonogenic assay performed at a lower dose (5Gy) to limit GBM cell death and chromosomal instability induced by higher doses. Fifteen R5CM and eighteen R5SASP clones were obtained from clonogenic assay performed in CM or SASP respectively. Molecular profile analyses by DGE-sequencing for each set of clones revealed a wider distribution in gene expression in R5SASP clones as compared to R5CM (*Supplementary Figure 2*), suggesting a stronger molecular diversity within these clones. To confirm this hypothesis, we calculated the coefficient of variation (CV) for all genes, which globally were significantly higher in R5SASP than in R5CM (*Figure 1g*). We further identified a set of 130 genes with a coefficient of variation>1 and a differential expression between CM and SASP >2. Within this set of genes, most of them were found in R5SASP set of clones (28 genes in R5CM and 91 genes in R5SASP; *Figure 1h*), strengthening that SASP increases the molecular diversity of R5SASP clones.

**Table 1:**
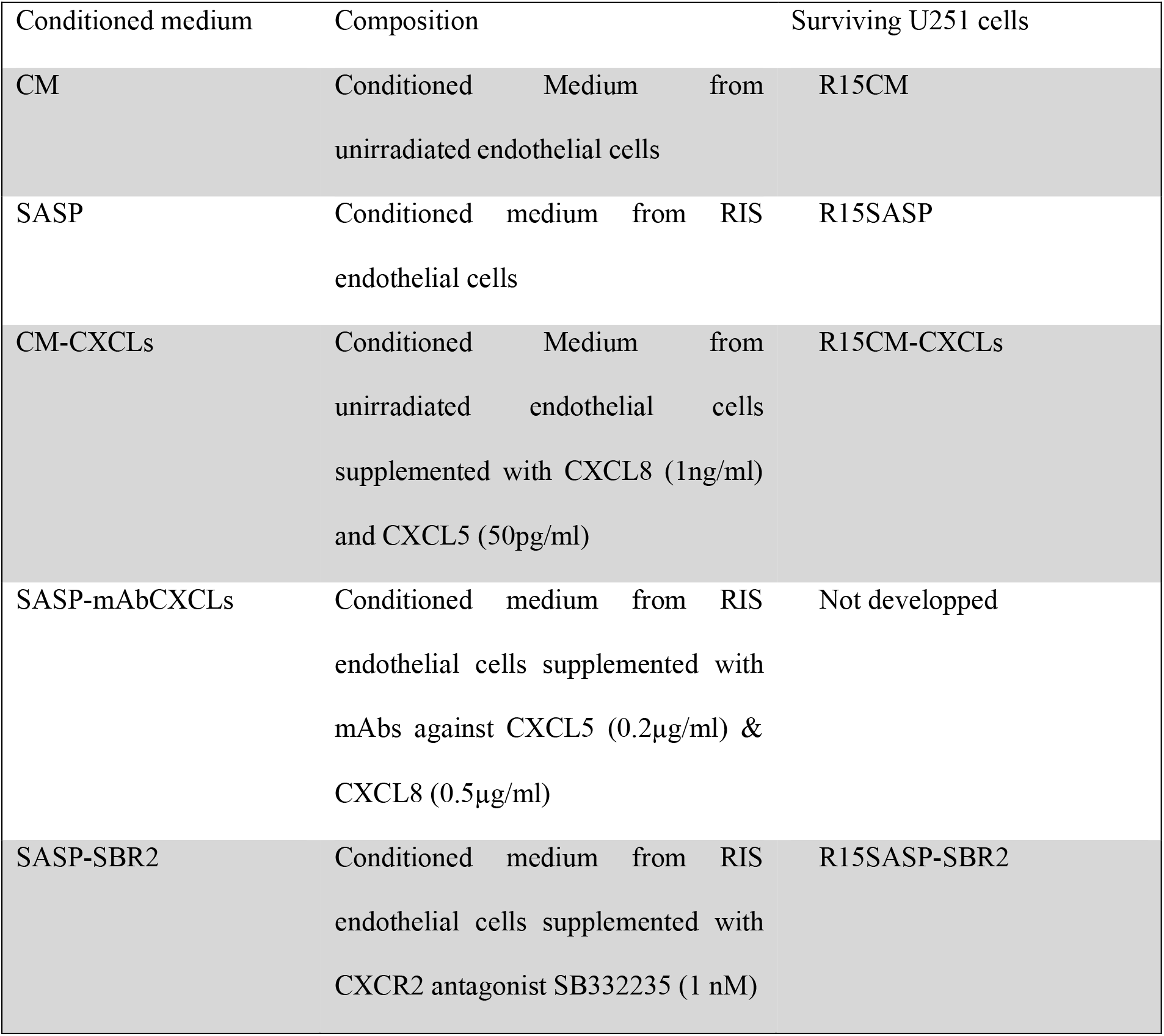
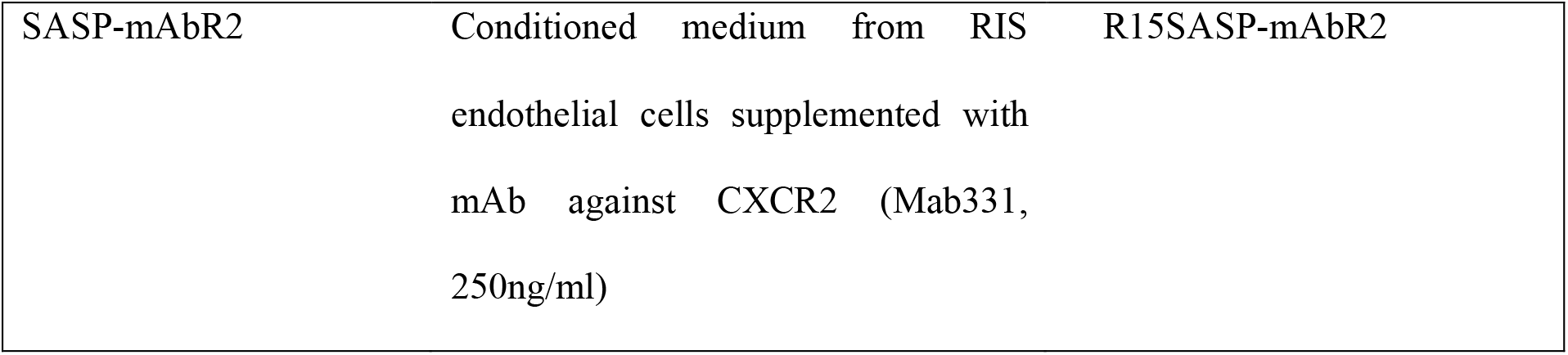
Nomenclature and characteristic of conditioned media from RIS endothelial cells and related 15Gy-radiation surviving U251 cells

**Figure 1:**
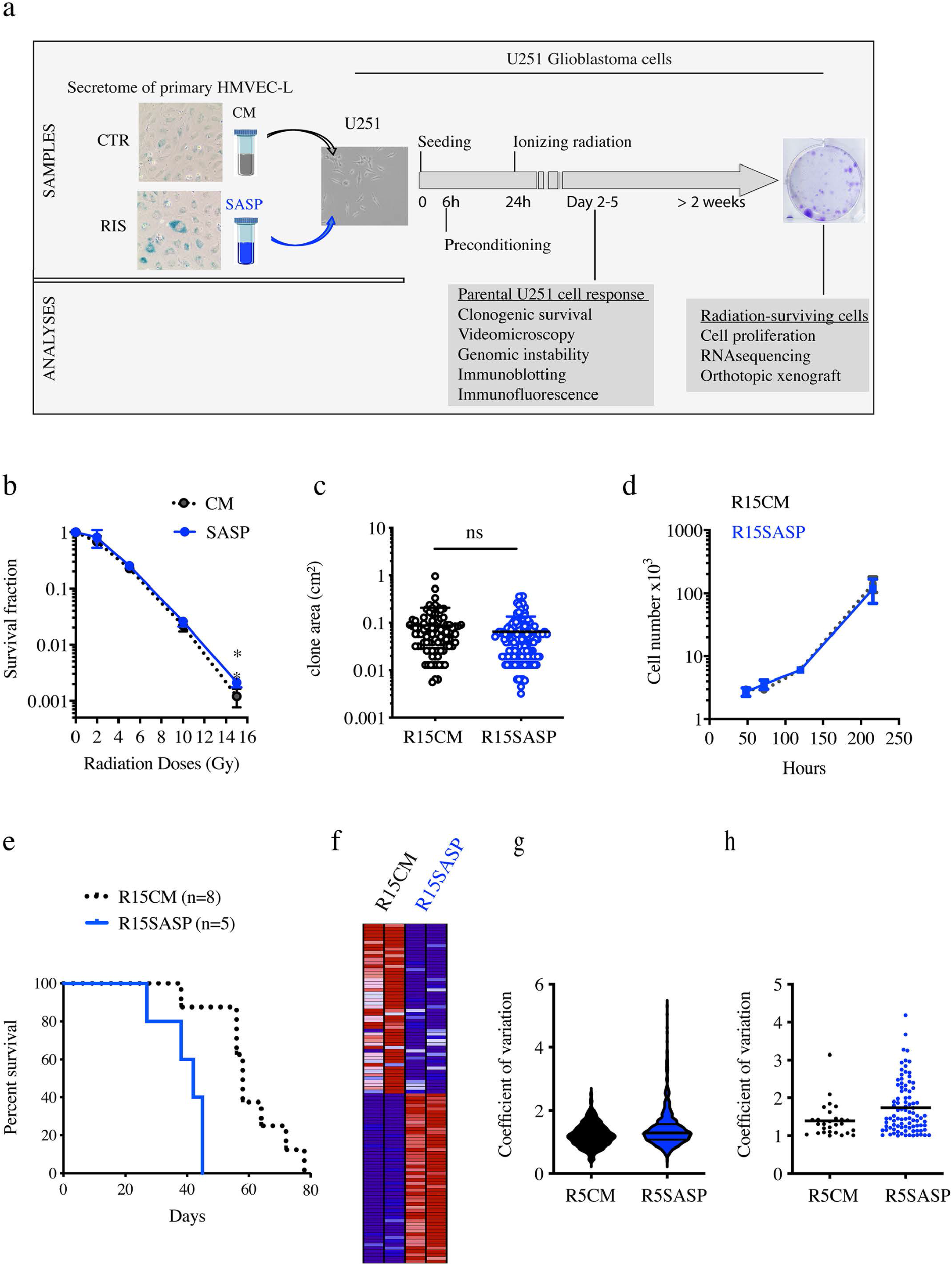
SASP increases aggressiveness of surviving GBM cells to radiation *in vivo*. a. Schematic plan of experiments. Secretomes (CM and SASP) were collected from primary endothelial HMVEC-L cells that were respectively unirradiated or after induction of 15Gy radio-induced senescence (RIS). glioblastoma U251 cells were plated at low density, preconditioned in CM or SASP after 6 hours, irradiated and cultured for 2 weeks. Cellular and molecular characterization after irradiation were performed on parental U251 cells and corresponding radiation-surviving cells. b. Radiation survival fraction of U251 cells depending on secretome preconditioning. U251 cells were preconditioned in CM (dashed black line) and SASP (solid blue line) prior to perform clonogenic assay in response to escalading doses of radiation (0 to 15Gy). After clone count, survival fraction was calculated. Results are presented as mean±SD (n≥3). Two-way ANOVA was performed for statistical analyses, * p<0.05. c. Size of R15CM (black) and R15SASP (blue) surviving clones following 15Gy-radiation. Each dot represents one clone. Mean±SD are also indicated (n=3). d. Proliferation of R15CM and R15SASP surviving clones. R15CM (dashed black line) and R15SASP (solid blue line) cells were collected and cultured in regular DMEM media. Cells were counted at 2, 3, 5 and 9 days. Results are presented as mean±SD (n=3). e. Kaplan-Meyer plot of overall survival of NSG mice following orthotopic injection of R15CM (dashed black line, n=8) or R15SASP (solid blue line, n=5) surviving cells. Log-rank test p<0.005. f. Transcriptomic analysis using DGE-sequencing of R5CM or R5SASP surviving clones. Overexpressed genes are indicated in blue whereas underexpressed genes are indicated in red. g. Coefficient of variation calculated for each gene in R5CM and R5SASP clones. h. Coefficient of variation of genes altered specifically by CM or SASP.

Altogether, our results show that SASP alters tumor cell evolution during radiation exposure leading to more heterogeneous and more aggressive radiation-surviving GBM cells.

### SASP enhances tumor genomic instability in response to radiation

Because SASP treatment led to more heterogeneous and more aggressive radiation-surviving GBM cells, the impact of SASP on GBM radiation response was investigated by videomicroscopy over 5 days. As expected, 5Gy irradiation decreased U251 cell proliferation cultured in CM (*Figure 2a*). When U251 cells were cultured in SASP, we observed a slight decrease in proliferation of unirradiated U251 cells while SASP dramatically decreased proliferation after 5Gy (*Figure 2a*). SASP also increased radio-induced cell death (*Supplementary Figure 3a*). Interestingly, SASP-inhibition of proliferation was concomitant with an increased occurrence of abnormal divisions in days following radiation (*Figure 2b*) with an enhanced ratio of cells dividing in three cells or more, or unable to complete mitosis (*Figure 2c*). Indeed, while U251 cells cultured in CM displayed 12 ± 7% abnormal mitosis after irradiation, more than 45 ± 11% of mitosis were abnormal when cells were cultured in SASP (p<0.01). As expected, percentage of abnormal mitosis without radiation was below 5% for both media (*Supplementary Figure 3b*). Since abnormal divisions lead to chromosome mis-segregation and aneuploidy, polynucleated cells were counted and were twice more numerous in SASP as compared to CM 72 hours after 5Gy (*Figure 2d*).

**Figure 2:**
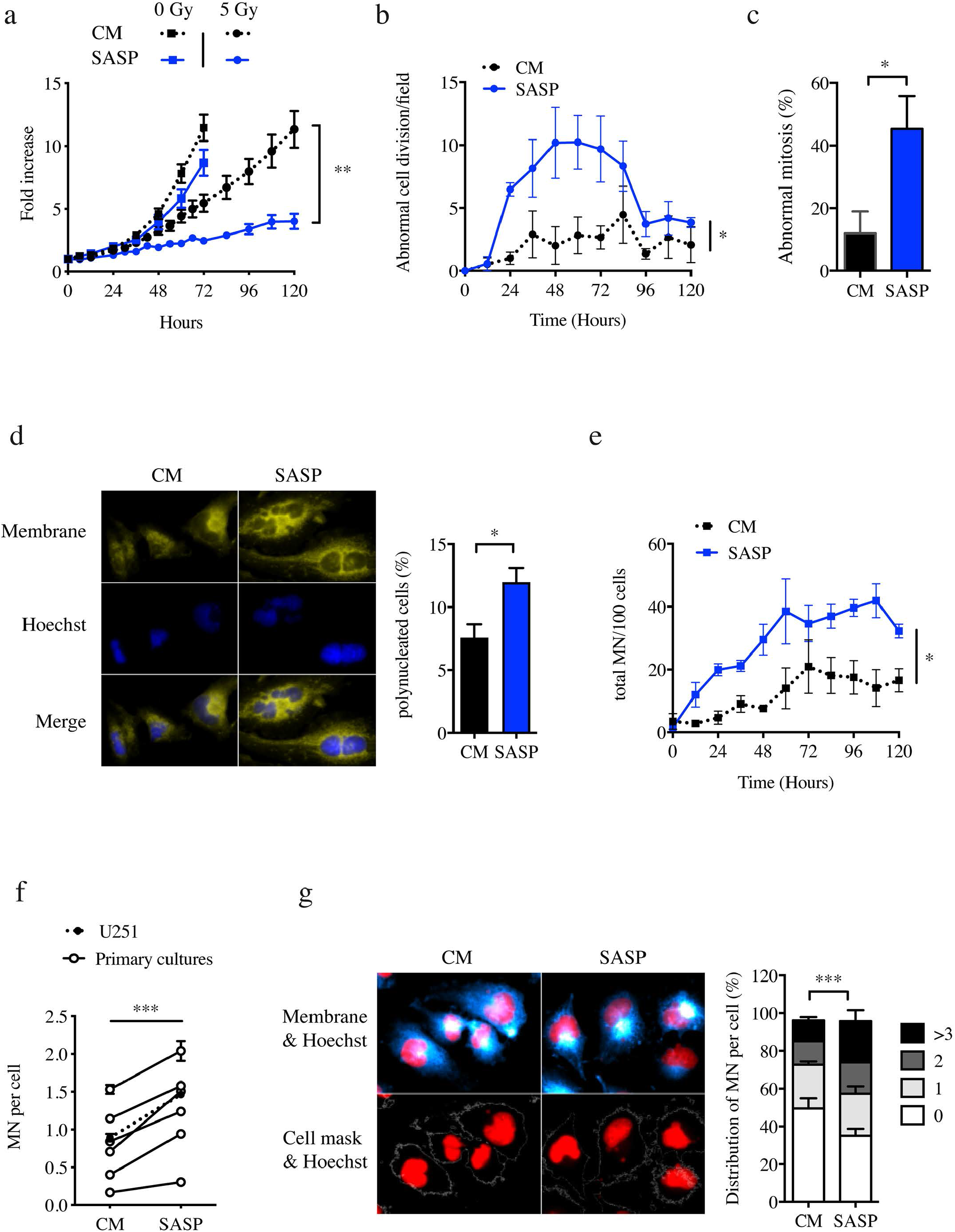
SASP enhances resistance and genomic instabilities after 5Gy-radiation. a. Monitoring of U251 cell proliferation by videomicroscopy analysis in CM (dashed black line) and SASP (solid blue line) in absence of radiation (squares) or after 5Gy-radiation (circles). Results are presented as the number of cells per field normalized to the initial number of cells (mean±sem, n≥3, Two-way ANOVA; ** p<0.01). b. Abnormal divisions of U251 cells in CM (dashed black line) and SASP (solid blue line) in response to 5Gy-radiation using videomicroscopy analysis. Abnormal divisions include cell division in more than 2 cells and failed mitosis. Results are presented as the relative frequency of abnormal cell division per field normalized to sham (CM and no irradiation) (mean±sem, n≥3, Two-way ANOVA; * p<0.05). c. Frequency of total number of abnormal mitosis as compared to total mitosis of U251 cells in CM (black) and SASP (blue) in response to 5Gy-radiation. (mean±SD, n=4, t-test * p>0.05). d. Frequency of polynucleated U251 cells in CM (black) and SASP (blue) 72 hours after 5Gy-irradiation. *Left panel*: representative pictures of membrane and Hoechst staining. *Right panel*: quantification of polynucleated U251 cells normalized to total number of cells (mean±sem, n=6, t-test; *p<0.05). e. Total MN (MN) production in U251 cells in CM (dashed black line) and SASP (solid blue line) in response to 5Gy-radiation using videomicroscopy analysis. Results are presented as the mean of MN in cells per field (mean±sem, n=3, two-way ANOVA; ** p<0.01). f. Micronulei (MN) formation in response to radiation in human primary GBM cells. Results are presented as the mean of MN per cell after 5Gy radiation in CM or SASP. Mean production of MN in U251 cells is indicated (black circle) (two-way ANOVA; *** p<0.001). g. Distribution of cells containing 0, 1, 2 or 3 MN. *Left panel*: representative pictures of membrane and Hoechst staining (top) and following macro analysis (bottom). *Right panel*: quantification of U251 cells with 0, 1, 2 and 3 MN 72 hours after 5Gy-irradiation (mean±SD, n=3, two-way ANOVA *** p<0.001).

Since abnormal mitosis and aneuploidy often result in micronuclei (MN) formation caused by mis-or unrepaired DNA damages, we counted the number of MN excluded from the nucleus during anaphasis. As expected, MN were only observed in response to radiation in a time dependent manner (*Figure 2e & Supplementary Figure 3c*). Interestingly, SASP triggered a faster and stronger production of MN after radiation both in U251 cells and in relevant models of human GBM primary cultures (*Figure 2e and 2f*). Besides this global increase in MN induction, the number of cells displaying 3 or more MN after 5Gy were two-fold higher in SASP than in CM (22% in SASP vs 11% in CM) (*Figure 2g*).

### SASP alters replicative stress and DNA elongation

To decipher the mechanisms behind SASP-induced genomic instability, we characterized the chromosomal composition and sequestration of DNA breaks in MN by FISH against centromere and telomere and *γ*H2Ax staining respectively. Whereas we previously reported that the number of MN per cell was significantly increased by SASP as compared to CM, their DNA compositions were similar in both conditions with essentially chromosome fragments lacking centromere and similar sequestration of DNA breaks (*Figures 3a, 3b*). Since damaged DNA enclosed in MN might be associated with failed DNA repair mechanisms and cell cycle checkpoints, stress-response pathways were analyzed in irradiated U251 cells in presence of CM or SASP. If no modulation were observed for most of the candidates involved in the reparasome, detoxification, cell cycle arrest, we observed an overexpression of the active form of CHK1, NPM and ERK1/2 after radiation in presence of SASP as compared to CM condition (*Figure 3c*). Specific pharmacological inhibition of each of these factors, respectively by NSC348884, LY2603618, and FR180204, prevented SASP-induced MN and reduced the number of polynucleated cell after irradiation (*Figure 3d and 3e*). Finally, we investigated whereas SASP impacts replicative stress by measuring replication fork progression. DNA fiber assays were performed after cell conditioning in CM or SASP 72 hours after 5Gy. As expected, when cells were cultured in CM medium, radiation shortened IdU track length in CM media (*Figure 3f*). Strikingly, when radiation was performed in SASP medium, the elongation rate was not reduced but rather significantly increased suggesting a dysregulation in this tightly regulated process.

In conclusion, SASP exacerbates radiation-induced genomic instability in GBM cells through several mechanisms involved in genome maintenance, including MN formation, DNA damage response machinery and DNA replication rate.

**Figure 3:**
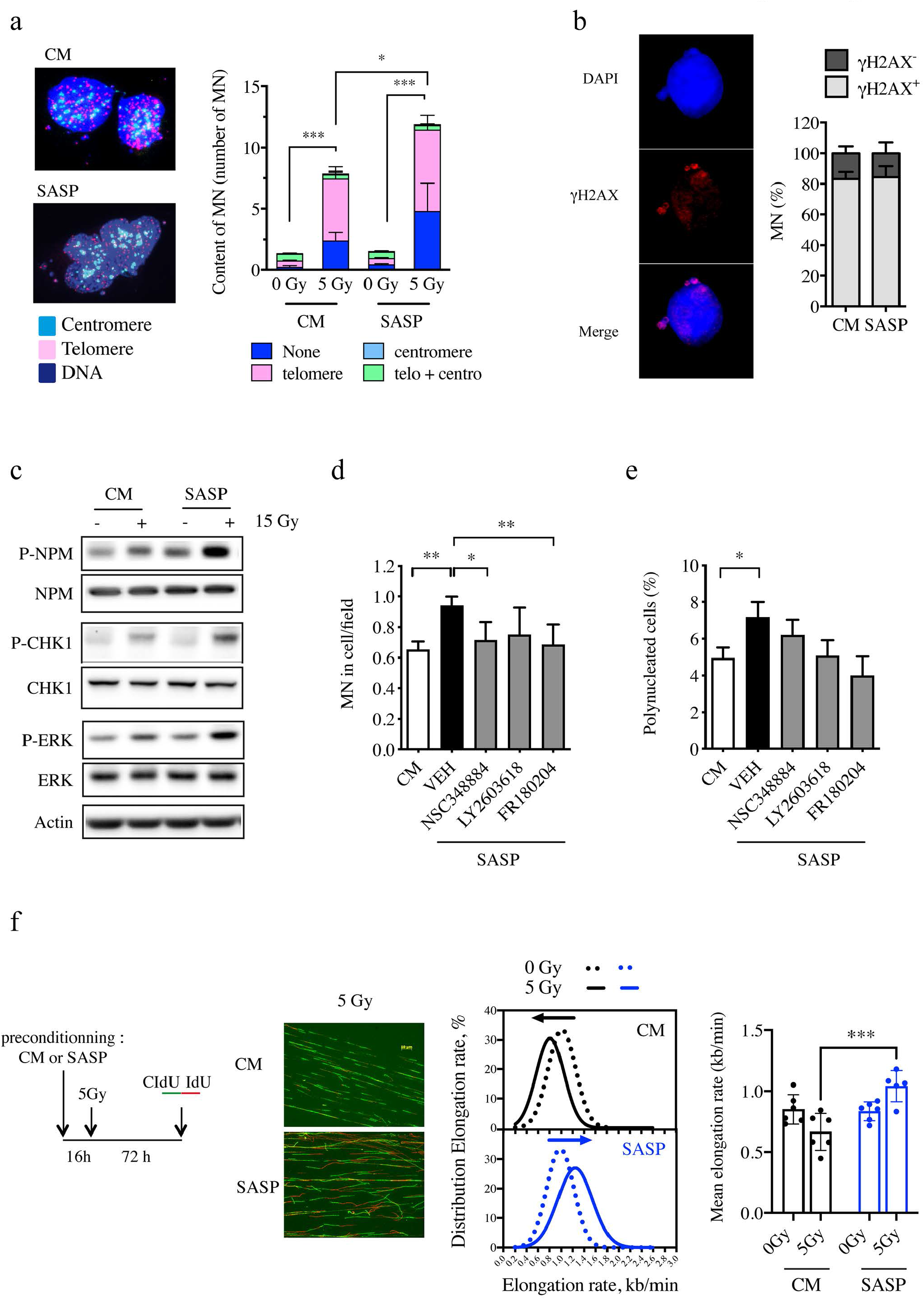
SASP alters replicative stress and DNA elongation after 5Gy radiation. a. Analysis of MN content using telomere and centromere probes. *Left panel*: representative pictures of immunofluorescence staining. *Right panel*: quantification of MN with centromere^+^ and/or telomere^+^ staining 72 hours after 5Gy-irradiation in CM or SASP. (n=3, two-way ANOVA; *p<0.05, *** p<0.001). b. Frequency of γH2AX positive MN in CM and SASP in response to radiation. *Left panel*: representative pictures of immunofluorescence staining. *Right panel*: quantification of MN with γH2AX^+^ staining 72 hours after 5Gy-irradiation. (mean±SD, n=3). c. Representative immunoblot of activation of NPM, CHK1 and ERK1/2 molecular pathways 24 hours in response to radiation in CM or SASP. Cells were irradiated at 15Gy. Actin was used as a loading control. (n=3). d. Production of MN in U251 cells after 5Gy-radiation in SASP in presence of NPM (NSC348884), CHK1 (LY2603618) and ERK1/2 (FR180204) inhibitors. e. Frequency of polynucleated cells U251 cells after 5Gy-radiation in SASP in presence of CHK1 (LY2603618), NPM (NSC348884) and ERK1/2 (FR180204) inhibitors. (mean±sem, n=6, one-way ANOVA, * p<0.05). f. Distribution of elongation rate in U251 cells in CM and SASP 72 hours after 5Gy-radiation. *Left panel*: scheme of experiment. *Median panel*: representative pictures of DNA fibers labeling. *Right panel*: distribution of elongation rate in U251 cells in CM (black) and SASP (blue) in Sham (dashed) and after 5Gy-radiation (solid). (mean±SD, n=3, two-way ANOVA; *p<0.05, *** p<0.001).

#### CXCL5 and CXCL8 trigger SASP-induced genomic instability in response to radiation

To identify the key factors involved in SASP-induced genomic instability, a molecular profiling was performed using RNA-sequencing analysis on unirradiated and RIS endothelial cells (*Figure 4a*). We identified 242 genes differentially expressed, the majority of them being up-regulated in RIS endothelial cells (*Figure 4a*). As expected, bioinformatics analyses showed among the top 10 enrichment pathways the modulation of angiogenesis or plasminogen activating cascade related to endothelial cell stress response and p53 transcriptional pathway involved in senescence induction (*Supplementary Figure 4*). Since the secretome of RIS endothelial cells specifically altered tumor cell genomic instability, further bioinformatic analyses focused on proteins involved in binding processes (*Supplementary Figure 4*) and 13 proteins were identified as potentially secreted factors (*Figure 4b*). We selected the two most up-regulated proteins, LIF and CXCL5, as well as two well-known cytokines secreted by endothelial cells, CXCL8 and IL-33. Whereas quantitative PCR confirmed an increase of all four genes expression, ELISA assays did not reveal significant changes of LIF and IL-33 secreted levels between CM and SASP (*Supplementary Figure 5a and 5b*). Remarkably, secreted amounts of both CXCL5 and CXCL8 were significantly increased in SASP as compared to CM (*Figure 4c*). Two other endothelial cell lines were analyzed for CXCL5 and CXCL8 expression in response to radiation, the human umbilical vein endothelial cell (HUVEC) and the immortalized human brain microvascular endothelial cells (HBMEC). An increase in mRNAs of both cytokines was observed in both HUVEC and HBMEC 10 days after radiation exposure (*Supplementary 5c*). The direct implication of CXCL5 and CXCL8 in GBM cell genomic instability was then validated either by blocking these two cytokines in SASP or by supplementing CM with both cytokines. First, blocking CXCL5 and CXCL8 with antibodies, either individually or in combination, prevented the formation of SASP-induced MN and polynucleated cells after irradiation (*Figure 4d*). Accordingly, addition of CXCL5 or CXCL8 in CM increased the production of MN and the number of polynucleated cells induced by irradiation (*Figure 4e*). Remarkably, the combination of both cytokines triggered the same formation of MN than the SASP itself. Finally, implication of CXCL5 and CXCL8 in NPM, CHK1 and Erk1/2 activation in response to radiation was investigated. Addition of each inhibitor in CM supplemented with CXCL5 and CXCL8 significantly reduced both MN formation and polynucleated cells frequency in response to radiation (*Figure 4f*).

**Figure 4:**
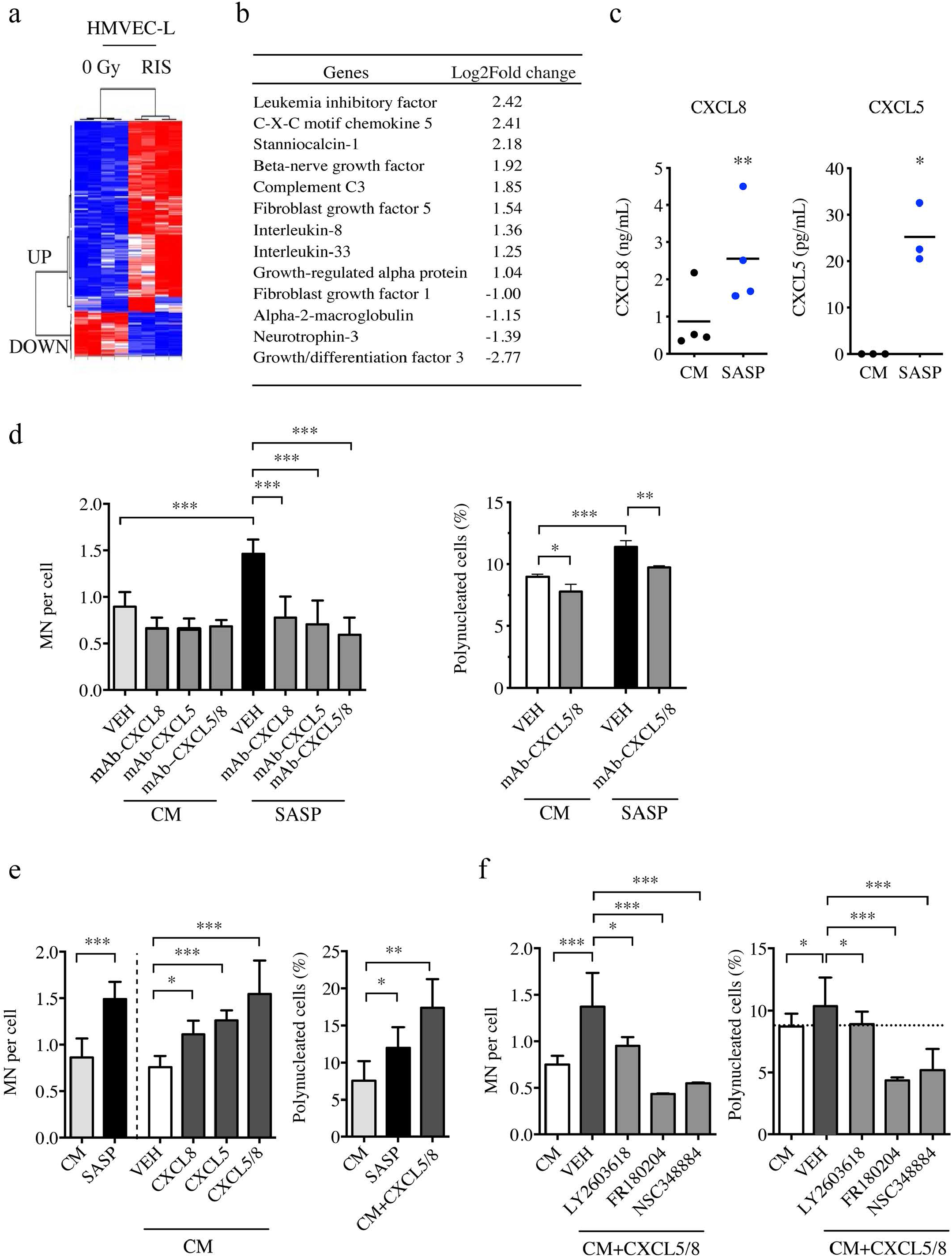
CXCL8 and CXCL5 secreted in SASP by EC increase radiation-induced genomic instability in GBM cells. a. Hierarchical clustering of the 242 genes differentially expressed in unirradiated (0 Gy) or radiation-induced senescent (RIS) HMVEC-L cells. Gene expression values are normalized using the mean of all samples and presented as a heat map (n=3, XLstat analysis). b. PANTHER gene analysis belonging to the “Receptor Binding” category with more than one log2 fold change in RIS EC compared to Sham EC. c. ELISA assay of CXCL8 and CXCL5 in CM and SASP. (n≥3, t-test; * p<0.05, ** p<0.01). d. Radio-induced MN production per cell *(left panel)* and polynucleated cell frequency *(right panel)* in U251 cells after 5Gy in CM or SASP supplemented with blocking antibodies against CXCL8 (0.5 μg/mL) and/or CXCL5 (0.2 μg/mL)(mean±SD, n=3, one-way ANOVA; ** p<0.01, *** p<0.001). e. Radio-induced MN production per cell *(left panel)* and polynucleated cell frequency *(right panel)* in U251 cells in CM, SASP or CM supplemented with CXCL8 (1 ng/mL) and/or CXCL5 (50 pg/mL) after 5Gy (mean±SD, n=3, one-way ANOVA; * p<0.05; ** p<0.01; *** p<0.001). f. Radio-induced MN production per cell *(left panel)* and polynucleated cell frequency *(right panel)* after 5Gy in U251 cells in CM, SASP or CM supplemented with a combination of CXCL8 and CXCL5 following addition of CHK1 (LY2603618), ERK1/2 (FR180204) and NPM (NSC348884) inhibitors (mean±SD, n=3, one-way ANOVA; * p<0.05; *** p<0.001).

#### SASP-induced genomic instability is transduced through CXCR2

CXCL5/8 signaling requires their binding to their receptor to mediate their signal. Both cytokines have been described to bind to the same receptor CXCR2, which is indeed expressed at the surface of GBM cells, both in U251 cell line and primary GBM cultures (*Figure 5a*). Interestingly, addition of either a CXCR2 antagonist (SB332235) or a specific monoclonal blocking antibody against CXCR2 (MAb331) reduced MN formation as well as polynucleated cells frequency in response to radiation (*Figure 5b and 5c*). Addition of CXCR2 blocking antibody did not alter the elongation rate of unirradiated U251 cells (*Figure 5d*). However, this process was significantly reduced by CXCR2 blocking antibody after irradiation in both CM and SASP. Thus, while blocking CXCR2 amplified DNA elongation stalling in GBM cells cultured in CM, it totally reverted the DNA hyper-replication driven by SASP after irradiation.

Altogether, these results showed that binding of CXCL5 and CXCL8 to CXCR2 fully recapitulates SASP-induced genomic instability in GBM cells in response to radiation and clearly establish CXCL5/8-CXCR2 axis as a key player in SASP-induced GBM instability.

**Figure 5:**
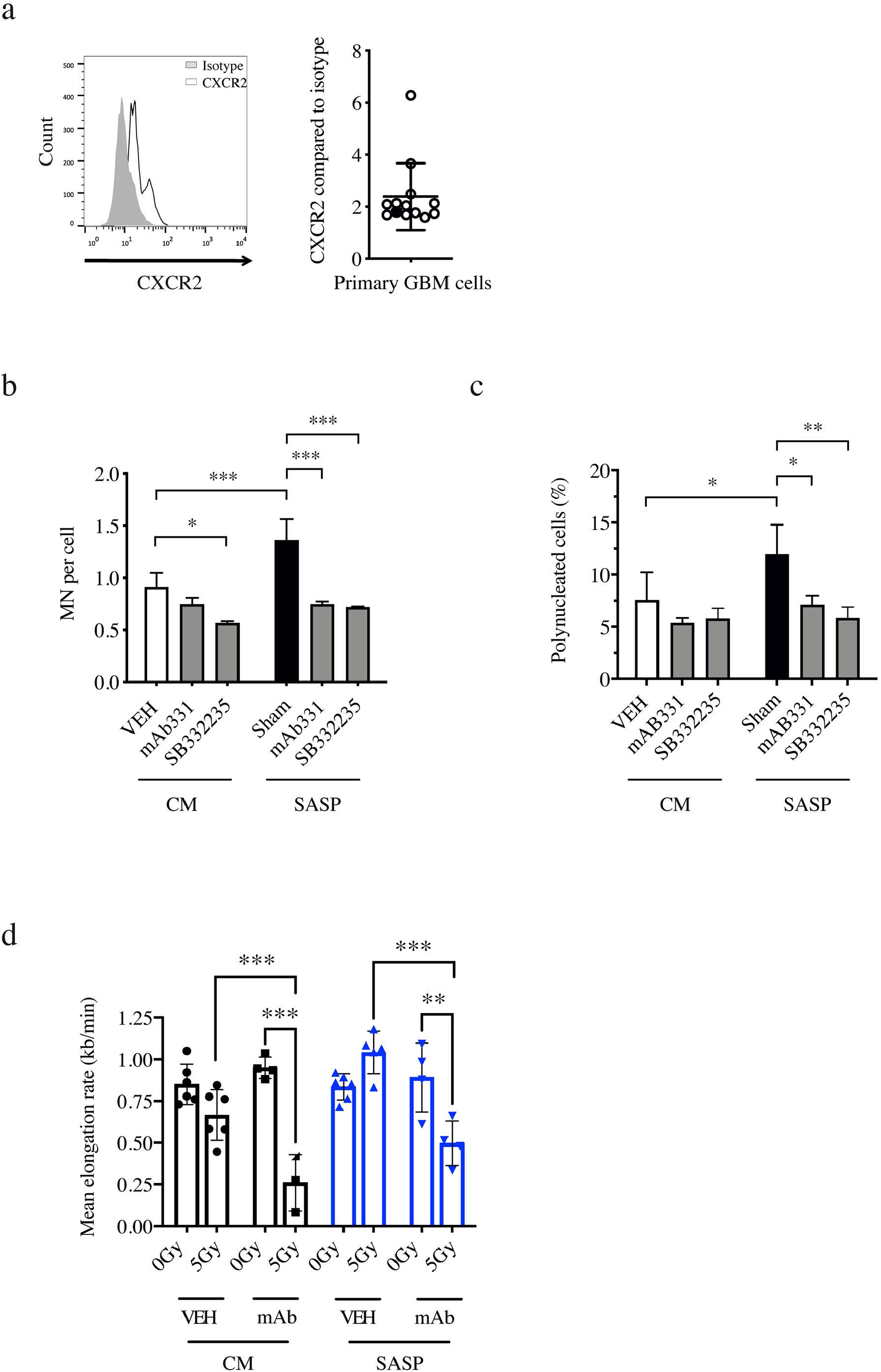
SASP increases radio-induced genomic instability through CXCR2. a. CXCR2 expression in U251 and human primary GBM cells by FACS analysis. *Left panel*: Representative histograms of CXCR2 staining in U251 cells. *Right panel*: Geomean of fluorescence compared to isotype. Geomean of CXCR2 expression in U251 cells and GBM primary cells is indicated respectively as black and white circles. b. Radio-induced MN production per cell after 5Gy in U251 cells in CM or SASP after CXCR2 blocking using monoclonal antibody MAb331(250ng/ml) or SB332235 antagonist (1nM). (mean±SD, n=3, one-way ANOVA; * p<0.05, *** p<0.001). c. Polynucleated cell frequency after 5Gy-radiation in U251 cells in CM or SASP after CXCR2 blocking using monoclonal antibody MAb331(250ng/ml) or SB332235 antagonist (1nM). (mean±SD; n=3; two-way ANOVA, * p<0.05, ** p<0.01). d. Elongation rate of U251 cells in CM (black) or SASP (blue) supplemented or not with CXCR2 monoclonal antibody (MAb331, 250ng/ml). When indicated, cells were irradiated with 5Gy-radiation. (mean±SD, n=3, two-way ANOVA; ** p<0.01, *** p<0.001).

#### CXCL5/8-CXCR2 axis mediates SASP-induced GBM aggressiveness in vivo

The relevance of the CXCL5/8-CXCR2 axis in SASP-induced GBM aggressiveness was analyzed in a murine orthotopic model of GBM. Clonogenic assays were performed to obtain 15Gy-surviving clones using U251 cells preconditioned and irradiated in either CM supplemented with CXCL5 and CXCL8, and SASP containing either CXCR2 blocking antibody or antagonist (*Supplementary Figure 6 & Table 1*). As previously described (*Figure 1*), 15Gy-surviving cells (namely R15CM-*CXCLs*, R15SASP-*MAbR2* and R15SASP-*SBR2*) were collected and injected into the cerebral subventricular zone of NSG mice. Interestingly, addition of CXCL5 and CXCL8 in CM led to the emergence of radiation-surviving cells as aggressive as the R15SASP (*Figure 6a and 6b, left panel*). Conversely, preventing CXCR2 activation in SASP, either with monoclonal antibody MA331b or with SB332235 antagonist, led to less aggressive tumor cells (*Figure 6a and 6b, middle and right panels*). These results clearly demonstrate that CXCL5/8-CXCR2 axis is involved on radiation-surviving GBM cell aggressiveness.

**Figure 6:**
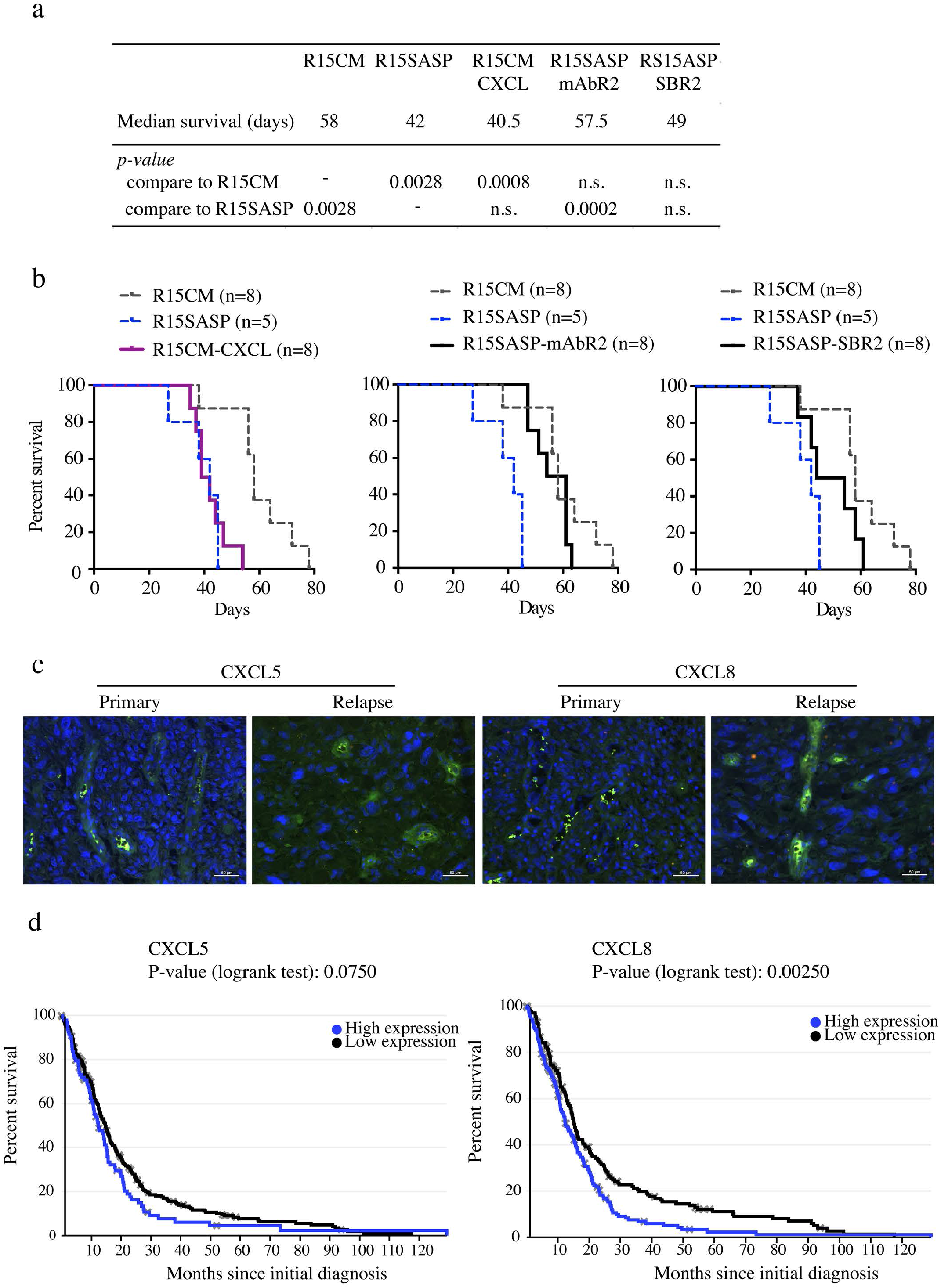
CXCL5/8-CXCR2 axis mediates SASP-induced GBM aggressiveness in vivo. a. Median survival (days) of tumor-bearing mice after orthotopic injections of radiation-surviving U251 cells obtained in presence of CM (R15CM), SASP (R15SASP), CM supplemented with CXCL8 and CXCL5 (R15CM*-CXCL5/8*), SASP supplemented with either CXCR2 monoclonal antibody Mab331 (R15SASP-*Mab*) or CXCR2 antagonist SB3322235 (R15SASP-*SB*). Log-rank p-values are indicated as compare to mice survival injected with either R15CM radioresistant cells or R15SASP radioresistant cells. b. Tumor-bearing mice survival after orthotopic injections of R15CM-CXCL (left panel), R15SASP-mAbR2 (middle panel) and R15SASP-SBR2 (right panel) (n=8/group) as compare to R15CM and R15SASP tumor-bearing mice. c. Immunofluorescence of CXCL5 and CXCL8 in primary and recurrent human GBM. d. Kaplan-Meier plots overall survival of GBM patients depending on CXCL5 and CXCL8 mRNA expression using the TCGA-glioblastoma database. (Log-rank tests analysis).

To determine the clinical relevance of our findings, we performed immunochemistry experiment against CXCL5 and CXCL8 on paired GBM biopsies, at diagnosis and relapse, from three patients. These patients had undergone the Stupp protocol, including radiation therapy, following GBM diagnosis. We did not detect CXCL5 and CXCL8 expression in initial primary GBM samples *(Figure 6c)*. Strikingly, staining of both CXCL5 and CXCL8 was observed around blood vessels in the GBM sample that recurred 123 days after the initial surgery. A lighter staining of both cytokines was observed in the second relapsing GBM that recurs 310 days after initial diagnosis. However, we were not able to detect CXCL5 or CXCL8 expression in the later relapsing GBM sample (recurrence at 327 days). To strengthen these results, gene survival association analyses were performed using TCGA database molecular profiling. Survival of GBM patient using the betastasis TCGA GBM databank for 299 patients revealed the worst overall survival for patients displaying the highest expression of either CXCL5 or CXCL8 (*Figure 6d*).

## Discussion

For decades, most strategies into improving outcomes from RT have been focused on overcoming radioresistance in tumor cells. This simple view has recently shifted towards recognizing the central importance of the TME ^15^ and a growing body of literature are now investigating the impacts of RT on TME and tumor resistance. At least, three recent publications reported that an irradiated TME increases GBM aggressiveness in orthotopic murine models ^6,16,17^ with no or limited investigation of underlying mechanisms. In the present work, using relevant *in vitro* and *in vivo* models, we demonstrate that alteration of the vascular TME by RT directly impacts relapsing tumor aggressiveness upon radiation. We identified the CXCL5/8-CXCR2 axis as a key driver altering genome maintenance of tumor cells and provided clinical relevance since expression of both chemokines was specifically detected in vessels from GBM relapse and positively correlated with worst patient outcome.

Genomic instability revealed by MN formation and aneuploidy is commonly observed during cancer initiation and response to treatment ^18,19^. Genotoxic therapies such as RT killed tumor cells trough accumulation of DNA damage. However, surviving cells undergo large genome remodeling following radio-induced genomic instability. RT induces accumulation of DNA damage and as a result cell cycle arrest and DNA replication fork stalling in order for the cell to repair DNA damage ^20^, as seen after 5Gy in GBM cells cultures in CM. Sustained replicative stress inducing aberrant replication fork structures has been previously linked to accumulation of DNA breaks ^21^ and genomic instability ^22^. In our study, we surely observed the opposite since SASP not only prevents radiation-induced replicative fork stalling but even triggers DNA hyper-replication. DNA hyper-replication has been reported in normal cells expressing activated RAS oncogene leading to accumulation of DNA damage and contributing to cell transformation ^23^. More recently, this process has been involved in the singular ability of mouse embryonic astrocyte to escape cellular senescence upon activation of RAS and elimination of RB and proposed to contribute to cancer incidence ^24^. Here, activation of CXCL5/8-CXCR2 signaling counteracts radiation-induced replicative stress and even induces a DNA hyper-replication. Interestingly, combination between SASP and RT activates at least, ERK, CHK1 and NPM, three members of stress response proteins, upstream of SASP-induced MN and aneuploidy. This overall molecular process impelling genomic instability causes the emergence of highly heterogenous radiation-surviving cells and increased tumor aggressiveness *in vivo*. Preclinical and clinical relevance of intratumoral heterogeneity in GBM aggressiveness has been demonstrated in several studies ^25–27^. In orthotopic GBM models, molecular heterogeneity resulting of various proportion of alternate molecular subtypes or diverse hybrid cellular states driven by the presence of GBM-stem-like cells decreased mice survival and limited therapy efficacy ^25,26,28,29^. In agreement with our model, recent single-cell analysis studies reporting that clonal heterogeneity appears as a highly dynamic process which does not derive only from the initial genetic diversity but also emerges as a fate decision by cancer cells driven by a beneficial outcome ^25,26,30,31^. Here, while we provided strong evidence on SASP-induced DNA hyper-replication, MN formation and aneuploidy, whether genomic instability involved stochastic or deterministic gene alterations has yet to be determined. For example, giving the reported role played by CXCL5/8 cytokines in tumorigenicity, GSC self-renewal and response to temozolomide through epigenetic alterations ^32^, whether SASP-induced genomic instability impacts tumor cell hybrid differentiation state should be investigated.

Numerous studies support the role of the microenvironment in shaping tumor phenotype toward a spatial and temporal heterogeneity. Through dynamic remodeling and cooptation, immune and stroma cells from the host provide a soil allowing the survival, progression and invasion of the tumor ^33^. Among the TME components, brain endothelial cells are described to create a unique perivascular niche that regulates GBM cell stemness, invasion and therapy resistance ^34^. Radiation induces early endothelial cell apoptosis enhancing cancer cell radiosensitization and favoring tumor growth control and regression ^35–37^. RT also triggers a well-described tumor bed effect characterized by an impaired neovascularization with reduced blood perfusion and low oxygen tension ^38^. This hypoxic microenvironment results in an extended latency period and a lower tumor growth rate, while enhances radiation resistance as compared to the same tumor budding in a more oxygenated bed ^39^. We recently reported that irradiation also induces endothelial cell senescence in the long-term ^14^. Interestingly, senescent endothelial cells are detected in 20Gy-irradiated brain of nude mice and in the tumor vicinity of post-mortem biopsies of recurrent glioblastoma from patients treated with 60Gy after surgery ^12,13^. In agreement with these results, we show expression of two secreted cytokines by RIS endothelial cells, CXCL5 and CXCL8, in GBM relapse biopsies around blood vessels. Brain endothelial cells are creating a perivascular niche where mutual GSC-EC interactions regulate GSC stemness, invasion and therapy resistance ^34^. These previous studies are limited to the impact of RIS endothelial cells to their ability to support GSC survival and growth. Here, we implement those results and demonstrate that RT chronically alters TME homeostasis creating a pro-tumorigenic vascular niche allowing the emergence of much more aggressive relapsing GBM cells. With the recent observation of senescent astrocytes in brain tissue from patients who have undergone brain radiation ^40^, we are not excluding the contribution of those cells, or other irradiated normal cells, in pro-tumorigenic niches. Thus, if forcing tumor cells to senescence by genotoxic stress has long been considered as an essential tumor suppressive mechanism that prevents tumor growth and dissemination ^41^, the chronic maintenance of a proinflammatory secretome by senescent cells also promotes tumor growth. Altogether, our results supported RT as a double-edged sword killing tumor cells and generating pro-tumorigenic senescent niches. Novel anti-cancer strategy must target those senescent niches ^41^, in order to maintain a homeostatic microenvironment and to retain the bright side of RT’s curative effects while getting rid of its dark side on tumor relapse. In this context, our study brings new insights in underlying molecular and cellular mechanisms involved in systematic GBM relapse and aggressiveness where radiation creates pro-tumorigenic niches of RIS endothelial cells that fuel radioresistant tumor cell emergence and aggressiveness. By deciphering this innovative concept, we can anticipate the development of next-generation targeted molecules, such as senotherapeutics, to limit GBM relapse and improve GBM outcome. Given the extensive use of RT in cancer treatments and relapse frequency, our study may help identify new approaches to treating cancer relapse not only in GBM but also in cancers with different origin.

## Methods

### Compounds

CXCL5 (#30-22) and CXCL8 (#200-08) cytokines were purchased from Peprotech. Blocking Abs against CXCL8 (Ab18672) and CXCL5 (Ab9802) were purchased from Abcam. Anti-CXCR2 blocking antibody (Mab331) and antagonist drug (SB332235) were purchased from R&D and TOCRIS (Bioscience) respectively. LY2603618, FR180204 and NSC 348884 inhibiting CHK1, ERK1/2 and NPM were respectively purchased from Bioscience, Selleckchem and from Axon Medchem.

### Cell culture and irradiation procedure

Primary human lung microvascular endothelial cells (HMVEC-L) were seeded at 5000 cells/cm^2^ in EBM-2 with 5% FBS and EGM-2 supplement (all products, Lonza Bioscience) until reaching complete confluence ^14^. Human Umbilical Vein Endothelial Cells (HUVEC) and immortalized human Brain Microvascular Endothelial cells (HBMEC) were seeded at 5000 cells/cm^2^ in EBM supplemented with 5% FBS and EGM supplements (all products, Lonza Bioscience) until reaching confluence ^42,43^. Human glioblastoma cell line U251-MG (ATCC) was cultured in DMEM 5 mmol/L glucose with 10% FBS, 1% Penicillin/Streptomycin and 2 mM Glutamine. Primary GBM cultures were obtained after mechanical dissociation from GBM patient biopsies in accordance with the ethical standards of the ethic national research committee and with the 1964 Helsinki Declaration and its later amendments or comparable ethical standards. These cells were cultured in Neural Stem Cell Culture Medium including DMEM/HAM-F12 supplemented with 2 mmol/L L-glutamine, N2 and B27 growth factors cocktails, 2 μg/mL heparin, 20 ng/mL EGF, 25 ng/mL bFGF, 100 U/mL penicillin and 100 μg/mL streptomycin, as previously described ^44^. All cells were grown in 5% CO2 incubators at 37°C, and monthly tested for Mycoplasma.

Cells were irradiated at a rate of 1.6Gy/minutes by CP160 (Faxitron) or by XRAD225Cx irradiator (Precision X-Ray) irradiators devices at different density in function of the experiments.

### Collection of HMVEC-L conditioned media and treatment to GBM cells

HMVEC-L were exposed to 0 or 15Gy once reaching complete confluence ^14^. Culture media were changed every week. Conditioned media were collected at day 21 and 28 when more than 30% of endothelial cells were senescent. Medium were centrifuged 5 minutes at 3000 rpm and then directly frozen in liquid nitrogen before storage at -80°C. For experiments, 6 hours after seeding of GBM cells in regular culture medium, equal amount of conditioned media (vol 1:1) were added. GBM cell preconditioning was performed for 18 hours before irradiation (Table 1).

### Radiation-surviving U251 clones

Radiation-surviving U251 cells are derived from U251-MG cells pretreated with conditioned media for 18 hours before irradiation at 15Gy. U251-MG were plated in 6-well plates at low density. 12 days after irradiation, the whole cell population was collected after trypsinization and grown in corresponding conditioned medium. Radiation-surviving cells were used at early passages following amplification for all experiments. Nomenclature of the different radiation-surviving populations are described in Table 1.

### Time-lapse assay

U251 cells seeded in 12-well-plates at 7500 cells/well were preconditioned 6 hours after seeding as described previously. Fourteen hours later, 0.5µM SiR-DNA (Spirochrome) were added to cellular medium to stain DNA. Cells were irradiated at 5Gy 4 hours later. Then, cells were followed by fluorescence videomicroscopy for 5 days (Eclipse Ti-E, Nikon). Brightfield and far-red (λabs/Em 652/674nm) images were taking every 10 minutes in 4 different fields per well. Number of cells, micronuclei and abnormal divisions were then quantified manually.

### Genomic instability

GBM cells were seeded in 12-well plates at 10000 cells/well, preconditioned and irradiated at 5Gy as described above. Cells were fixed in 4% paraformaldehyde 72 hours after irradiation and stained for DNA with Hoechst33342 for 10 minutes at room temperature. Ten pictures per well representing at least 100 cells were captured by fluorescence microscopy (λabs/Em 350/461nm, 60X magnification Zeiss). Mean number of micronuclei (MN) per cell was manually quantified from at least 100 cells, under open source software Fiji. MN were recognized by distinct Hoechst-positive vesicles filled with DNA fragments surrounding the nucleus. For primary GBM cultures, only adherent cell cultures were selected for the experiment. To count the number of MN and nuclei per cell (polyploidy), cells were counterstained with Hoechst33342 and 0.5X Deep Red Plasma membrane stain (Thermo) to outline cell surface. Ten pictures per well were captured by fluorescence microscopy (60X magnification Zeiss). Mask was applied using Fiji software to delineate cells. Number of MN per cell and polynucleated cell were manually counted for each field for a least 100 cells per condition.

### Fluorescent immunostaining

GBM cells were seeded in 12-well plates at 10000cells/well on glass coverslip coated with 1% gelatin and then preconditioned and irradiated at 5Gy in conditioned medium as described above. Cells were fixed in 4% paraformaldehyde for 10 minutes, permeabilized in 0.1% Triton X-100 for 10 minutes and blocked in PBS-5% goat serum for 30 minutes. Cells were incubated with primary antibody 1/200 γH2AX (9718, Cell Signaling) overnight at 4°C and secondary antibody 1/200 anti-IgG-Alexa568 (A11036, Thermofisher) during 1 hour at room temperature. Coverslips were mounted on slides with Prolong Gold with DAPI as DNA counterstain (P39635, Life technology), then observed on laser confocal microscope (model FV1000, Olympus or Nikon-A1). Images were processed using Image J.

### TC-FISH

GBM cells were seeded in 6-well plates at 10000cells/well and then preconditioned and irradiated in conditioned as described above. Metaphase preparations and Q-FISH were performed 72 hours later as described in ^45^. Briefly, cells were fixed with 4% formaldehyde for 2 minutes at room temperature, then treated with pepsin (0.5mg/ml) for 7minutes at 37°C, fixed again in 4% formaldehyde for 2 minutes, and finally sequentially dehydrated by 50%, 70%, 100% ethanol before being air-dried. The slides incubated with 50μl probe solution (0.3μg/ml for telomere and centromere) were denatured at 80 °C for 3minutes and incubated in the dark for 1 hour at room temperature. Genomic and MN DNA were respectively hybridized with a Cy-3-labeled PNA probe for TTAGGG sequence and a FITC-labeled PNA probe specific for centromere sequences (both Panagene). After hybridization, slides were washed 3 times with 70% formamide/10mM Tris pH 7.2, for 15 minutes and then with 50mM Tris pH 7.2/150 mM NaCl pH 7.5/0.05% Tween 20 for 5 minutes. Cells were counterstained with DAPI and mounted with PPD (1 mg/mL p-phenylenediamine-90% glycerol-10% PBS). Images were acquired by AutoCapt software (MetaSystems). Images of metaphase cells were acquired with a charge-coupled device camera (Zeiss) coupled with a Zeiss Axioplan microscope. Quantification image acquisition and analysis were performed using Isis software (version 3.9.1, MetaSystems). MN assays and Telomere and centromere staining are described step by step in https://www.youtube.com/watch?v=gc5uyTyHTrU&t=49s and https://www.youtube.com/watch?v=RqI1ulPWD_E.

### Clonogenic assay

Tumor cells were seeded in 6-well plates accordingly to their respective plating efficiency (PE), pre-conditioned and irradiated with escalating doses from 0 to 15Gy as described above. Twelve days after irradiation, isolated colonies were manually counted after staining with crystal violet (Sigma-Aldrich). PE and survival fraction (SF) were calculated using the following formula:

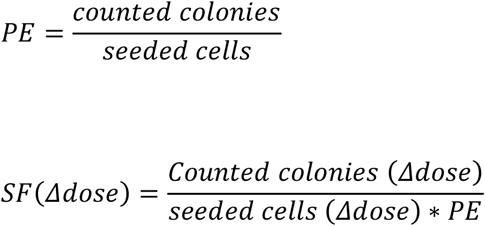

### ELISA assay

LIF (BMS #242, Invitrogen), IL-33 (BMS #2048, Invitrogen), CXCL8 (#, R&D systems) and CXCL5 (EHCXCL5, Invitrogen) quantification in endothelial media were performed by ELISA on HMVEC-L conditioned media according to the manufacturer instruction.

### Transcriptomic analysis

RNA extraction, quantity and quality evaluation were processed as described in ^44^. RNAs were extracted after 15Gy irradiation on day 21 for HMVEC-L, and day 10 for HUVEC and hBMEC. RNA from radiation-surviving U251 cells were extracted following their amplification in U251 medium for 2 weeks. Reverse transcription and qPCR were performed as described previously ^44^. Primer sequences for targeted genes are given in the Table 2. HGPRT, TATA and GAPDH were used as housekeeping genes.

**Table 2:**
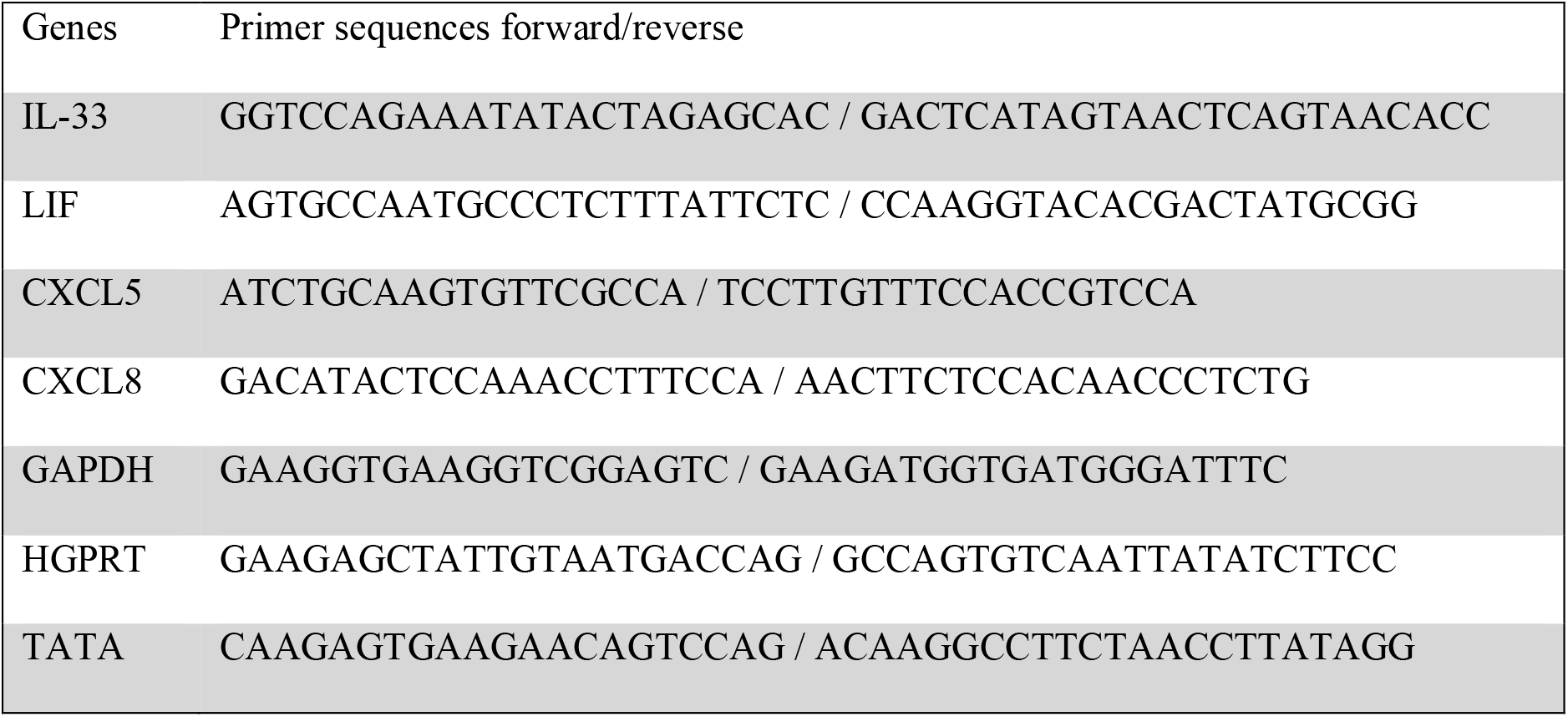
Primer sequences of genes of interest

For RNAsequencing using DGEseq, after demultiplexing and quality control with fastQC_0.11.2 (http://www.bioinformatics.babraham.ac.uk/projects/fastqc/), illumina adapters were trimmed with cutadapt-1.2.15 and reads with Phred. Quality score below 30 were filtered with prinseq-lite-0.20.36. Reads were aligned against human hg19 reference genome with tophat2.0.107. Reads count was realized with htseq-count from HTSeq-0.5.4p58 and differential analysis with DESeq2. Data were further analyzed using String software (version-10-5), a protein interaction database generating network of proteins interactions with a confidence cut-off of 0.6.

### Transcriptomic heterogeneity

5Gy-surviving U251 clones (R5CM and R5SASP) were obtained by clonogenic assay as described above, except that isolated clones were collected and cultured individually in corresponding conditioned medium, CM and SASP. Nomenclature of radiation-surviving clones were called R5CM and R5SASP followed by clone number depending on the preconditioning medium. Fifteen R5CM and eighteen R5SASP clones were collected. RNA extraction and DGEseq analysis were performed for each individual clone as described above. For each set of clones (R5CM and R5SASP), total number of reads as well as the mean (µread) and standard deviation (dread) for each gene were determined to measure the transcriptomic diversity (*Supplementary Figure 2*). Genes with µread<1 for all clones were excluded of further analysis. Then, the coefficient of variance (CV) representing clones scattering was calculated for each gene using the following formula 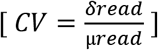. To address specific gene variability induced by the conditioned media, we selected genes with high variance (CV>1) and significant differential expression between R5CM and R5SASP (fold change >2).

### Flow cytometry

CXCR2 expression in GBM cells were assessed by flow cytometry using FACSCalibur (BD bioscience). Cells were incubated with CXCR2 mAb (555933 BD Pharmingen) for 30 minutes. Geomean of fluorescence from 10000 live events using CellQuest software (BD bioscience) were analyzed with FlowJo Software (FlowJo LLC, USA).

### Western blot

Western blots were performed as described in ^36^. Cells were lysed in Lysis buffer (10 mM Tris-HCl pH 7.5, 150 mM NaCl, 1% NP-40, 5 mM EDTA, 1 mM PMSF, 2 mM Na3VO4, 1 mM NaF, 0.2 μM Leupeptin, 0.3 μM Aprotinin). After denaturation, 20µg of protein lysates were loaded and run onto SDS-PAGE gels, then transferred to PVDF Immobilon-P membranes (Millipore). After blocking, membranes were incubated overnight at 4°C with the following primary antibodies: phospho-Thr202/Tyr204 Erk1/2, phospho-Thr199 NPM, phospho-Ser345 CHK1 (respectively #4370, #3541, #2348; Cell Signaling) and actin (05-636, MAB1501; Millipore). Following incubation for 1 hour at room temperature with HRP-coupled secondary antibodies (Jackson ImmunoResearch), proteins of interest were revealed by Clarity Western Blot ECL substrate (Millipore).

### GBM biopsies and Immunostaining

Three patients showing the following were selected from the french glioblastoma biobank (FGB, biobank number BB-0033-00093; https://www.linkedin.com/in/french-glioblastoma-biobank-808508153), with common criteria: 1) they underwent craniotomy for resection of primary IDH-wildtype GB; 2) they received the Stupp protocol, 3) They underwent reoperation for local tumor recurrence. All patients provided written informed consent according to FGB protocols and regulations ^46^. The delay between the 1st surgery and the recurrence was 124, 310 and 327 days.

Biopsy sections were deparaffinized and antigens were retrieved by boiling in 10mM sodium citrate buffer pH6 for 40 minutes followed by endogenous biotin blocking (Agilent) and normal goat serum blocking (Sigma-Aldrich). Sections were incubated overnight at 4°C with isotype controls and primary antibodies against human CXCL5 and human CXCL8 (Abcam), then detected with biotinylated secondary antibodies (Vector) and streptavidin FITC (Interchim). Nuclei were counterstained with DAPI (Sigma). Sections were observed under a fluorescence Axioscope 2 optical microscope (Zeiss) at x200 magnification. Pictures were taken from 5 randomly selected fields using 2 emission channels (λabs 365 and 488) and analyzed with Zen image-analysis system (Zeiss). Imaging in the 547-red channel was also done to counteract the green fluorescent background.

### DNA Fiber assay

GBM cells were seeded in 12-well plates at 10000cells/well and preconditioned as described above. Cells were irradiated at 5Gy 18 hours after addition of conditioned media. Cells were sequentially labeled with 25 μM CldU and 250 μM IdU (both Sigma-Aldrich), and DNA fiber spreads were prepared as described ^47^. Briefly, DNA fiber spreads were prepared from 0.5 × 10^6^ cells/ml. Slides were incubated in 2.5M HCl for 90 minutes, and then incubated in blocking buffer (3% BSA, 0.1% Tween in PBS) for 1 hour. Acid-treated fiber spreads were incubated with 1/1000 rat anti-BrdU mAb (Bio-Rad) for 1.5 hour to detect CldU, followed by 1/1500 mouse anti-BrdU mAb (BD) overnight to detect IdU. Then, 1/500 goat anti-rat AlexaFluor555 and 1/500 goat anti-mouse AlexaFluor488 (both Millipore) were then incubated for 1.5 hour. Slides were mounted in Immuno-Fluor mounting medium (MP Biomedicals). Fiber tracts were examined using an AxioVert 200 M fluorescence microscope (Zeiss). Pictures were taken from randomly selected fields with untangled fibers and analyzed with Fiji software. For fork speed analyses, lengths of CldU and IdU tracks were measured and converted into DNA bases (1 μm = 2.59kb) ^48^. Replicon clusters are stable units of chromosome structure, evidence that nuclear organization contributes to the efficient activation and propagation of S phase in human cells (Jackson and Pombo, 1998). A minimum of 100 individual fibers was analyzed for each experiment.

### Orthotopic injections of U251 human tumor cells in NSG mice

Immunodeficient male NOD.Cg-PrkdcscidIl2rgtm1Wjl/SzJ (NSG, Charles River Lab) mice were housed in animal facility (SFR F. Bonamy Inserm UMS026, Nantes) approved by the French Association for Accreditation of Laboratory Animal Care (AFSTAL) and maintained in accordance with the regulations and standards of Inserm and French Department of Agriculture. Orthotopic injections of radiation-surviving U251 cells (10^4^ cells in 2 μl PBS) were performed in the subventricular zone of brain (2mm on the right of the medial suture and 0.5mm in front of the bregma, 2.5mm depth:) of 8–12-week-old mice using a stereotaxic frame (Stoelting) as previously described ^49^. Animals were observed daily and euthanized after occurrence of clinical morbidity symptoms, mainly reduced mobility or more than 10% of weight loss.

### Statistical analysis

Experiments were performed at least three times and all statistics were investigated by Prism 7.0 (GraphPad Software). Student’s or ANOVA tests with 95% confidence estimation were performed on molecular or cellular experiments and Mantel log-rank test was used to animal survival. p value of less than 0.05 was taken to indicate statistical significance: * p<0.05, ** p<0.01, *** p<0.001.

## Supporting information

all sup data

## Acknowledgements

We thank the microscopy platform MicroPICell, the cytometry and the genomic core facilities Cytocell and BIRD as well as the animal facility UTE. We thank Julie Gavard for her gift of hBMEC. We thank La Ligue contre le Cancer, la Fondation ARC, the region Pays de la Loire and the SIRIC ILIAD for their financing support (ERRATA project).

We also thank the French Glioblastoma Biobank, in particular Pr Philippe Menei, Gwénaëlle Soulard (CHU, Angers), Dr Odile Blanchet and Mélanie Flipeau (CRB, CHU, Angers) and Pr Audrey Rousseau (Laboratoire d’Anatomie Pathologique, CHU, Angers).

