## Supplementary material for "Mechanistic insights of radiation-induced endothelial senescence impelling glioblastoma genomic instability at relapse": all sup data

**Supplementary Figure**

**Supplementary Figure 1 Characterization of U251 cells preconditioned in CM and SASP.** a. Plating efficiency of U251 cells in CM (black) and SASP (blue). (mean±SD, n=3). b. Predicted protein-protein interaction network of differential proteins between R15CM and R15SASP data.

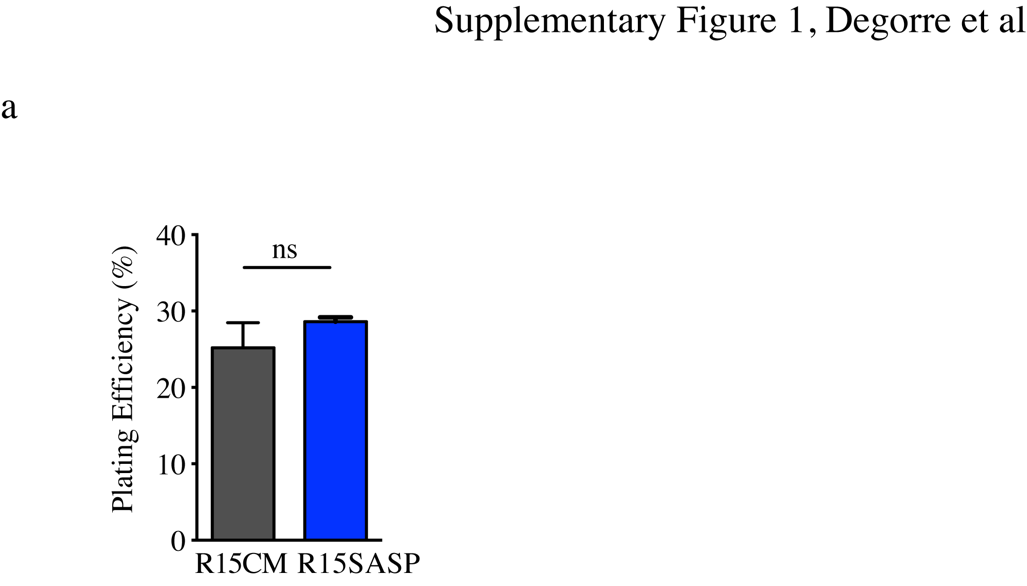

**Supplementary Figure 2 Molecular diversity of radiation-surviving clones depending on preconditioning medium.**

a. Means of total reads number of R5CM and R5SASP clones. b. Reads number per gene of R5CM and R5SASP clones. c. Coefficient of variation of all genes in R5CM and R5SASP clones.

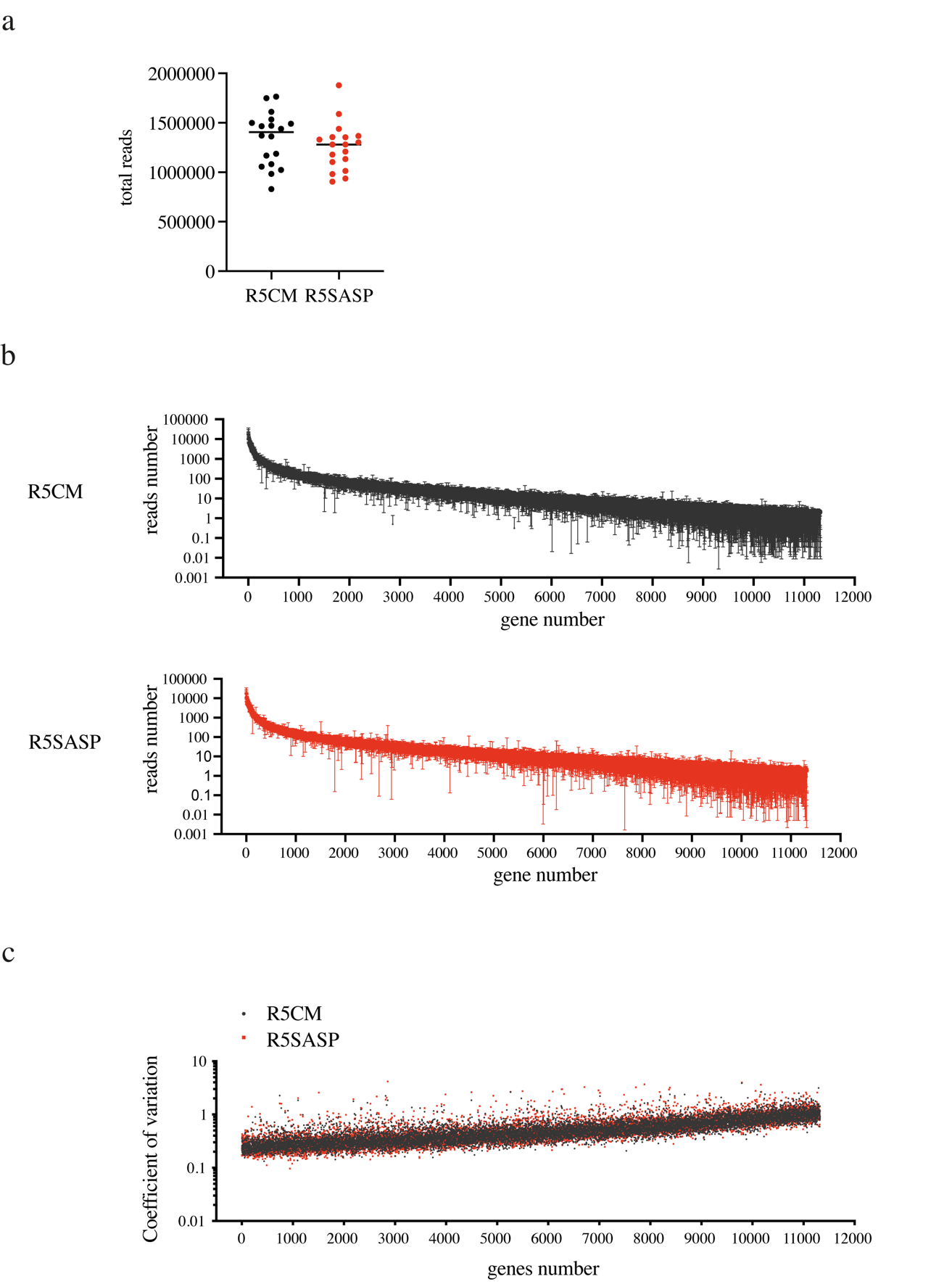

**Supplementary Figure 3 Cell death and MN formation in U251 cells.**

a. Dying U251 cells in CM (black) and SASP (blue) in response to 5Gy-radiation using videomicroscopy analysis. Results are presented as the mean of detached cells as compare to total number of cells per field (mean±sem, n=3). b. Frequency of total number of abnormal mitosis as compared to total mitosis of U251 cells in CM (black) and SASP (blue). (mean±SD, n=2). c. MN production in unirradiated U251 cells in CM (black) and SASP (blue) using videomicroscopy analysis. (mean±SD, n=2).

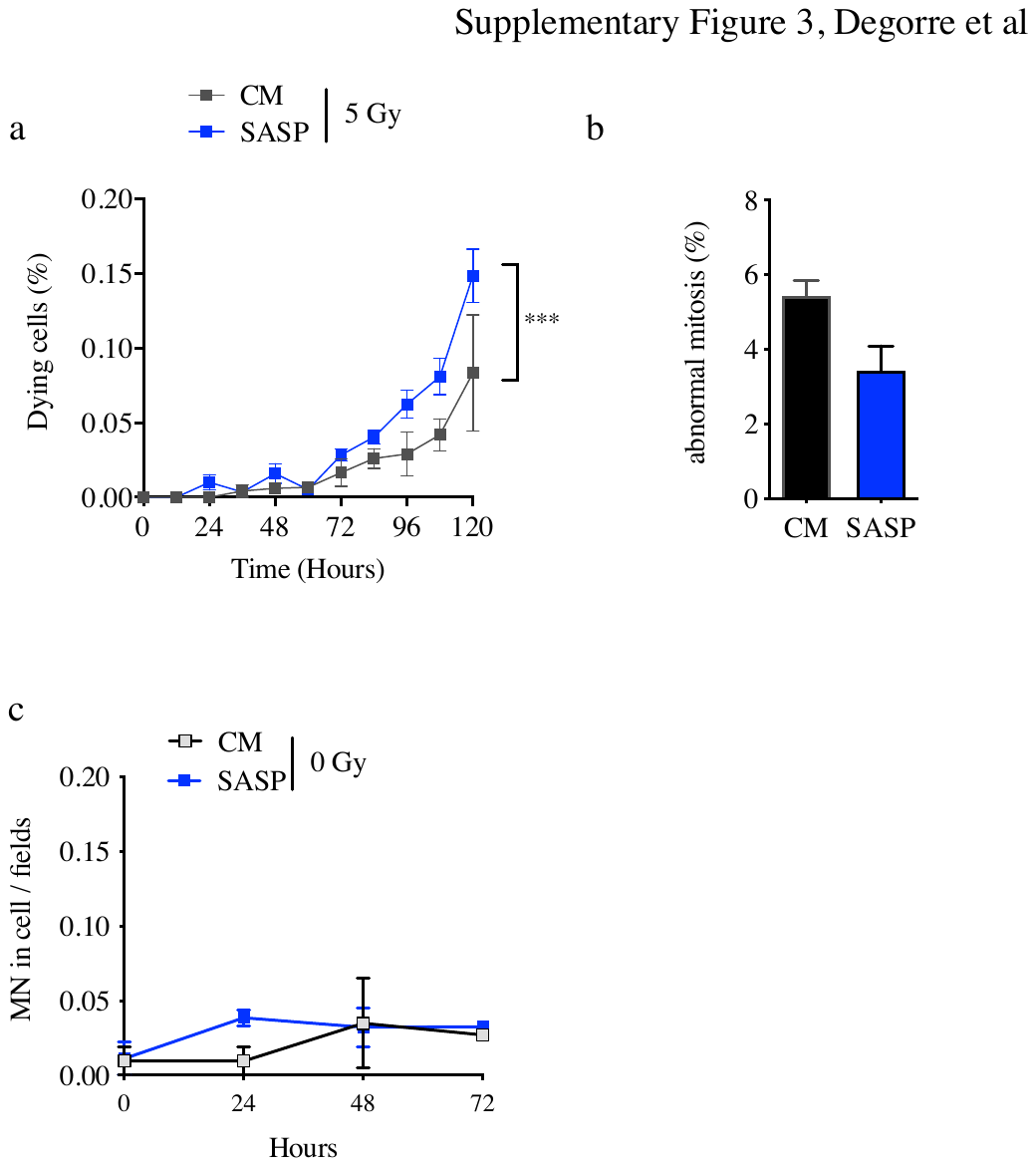

**Supplementary Figure 4 Transcriptomic analyses of the 10 most up-regulated pathways using PANTHER analysis.**

a: functional annotations of differentially expressed pathways are indicated (GO: Gene Ontology). b: transcriptomic analyses using PANTHER software for molecular functions (*left panel*) focusing on genes involved in protein binding (*middle and right panels*). Results are presented as pie charts.

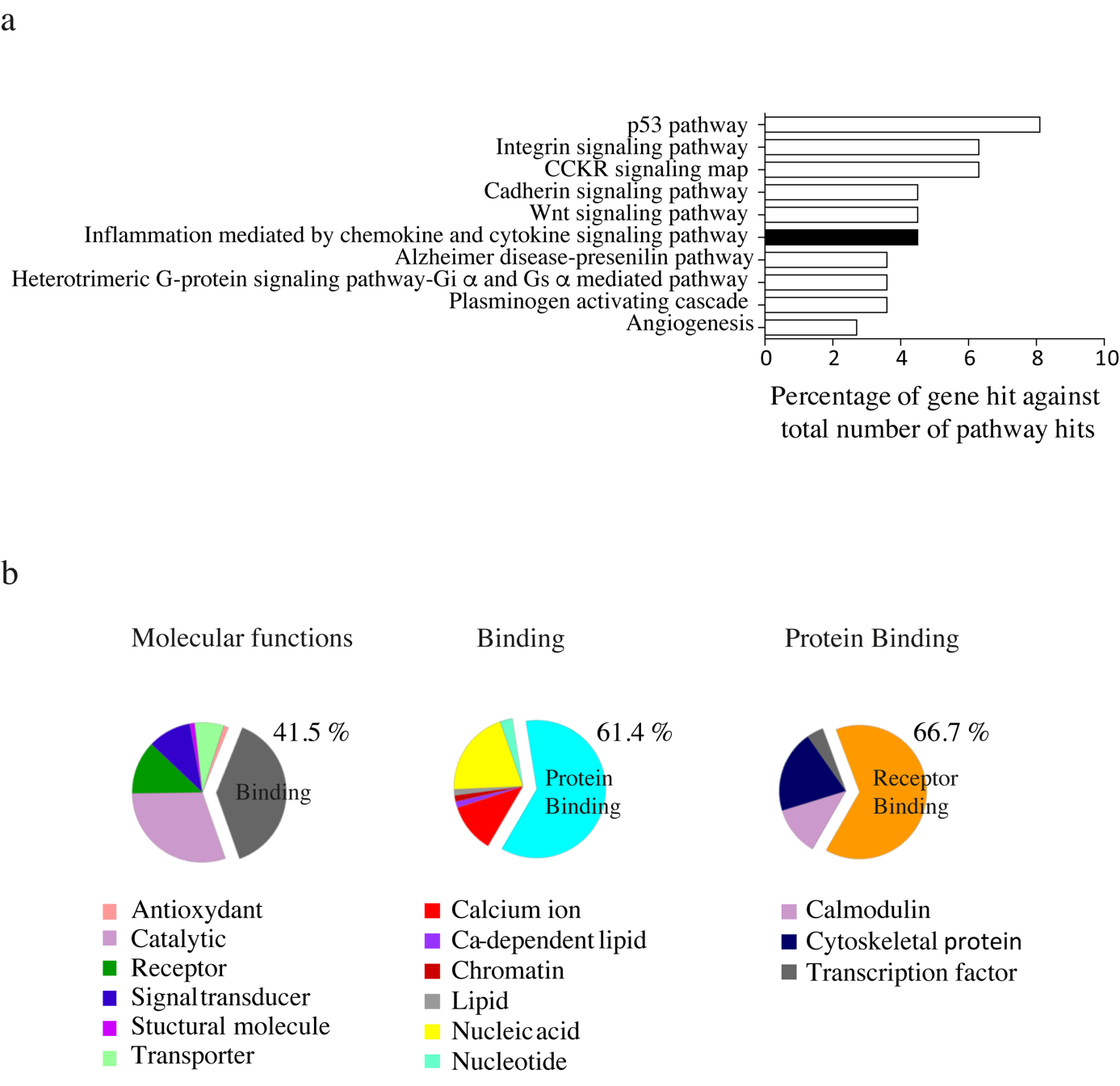

**Supplementary Figure 5 Cytokine expression in EC after 15Gy-radiation**.

a. Relative mRNA expression of IL-33, LIF, CXCL8 and CXCL5 in unirradiated (CTR) and radio-induced senescent (RIS) HMVEC-L (mean ± SD, n=3-4, paired t-test, * p<0.05, ** p<0.01, *** p<0.001). b. ELISA assay of IL-33 and LIF in CM and SASP from HMVEC-L 72 hours after 5Gy- radiation. (n≥3). c. Relative mRNA expression of CXCL8 and CXCL5 in human endothelial cell line HUVEC (n=2) and immortalized brain endothelial cells HBMEC (n=1) in absence of radiation (black) or after 15Gy-radiation (blue).

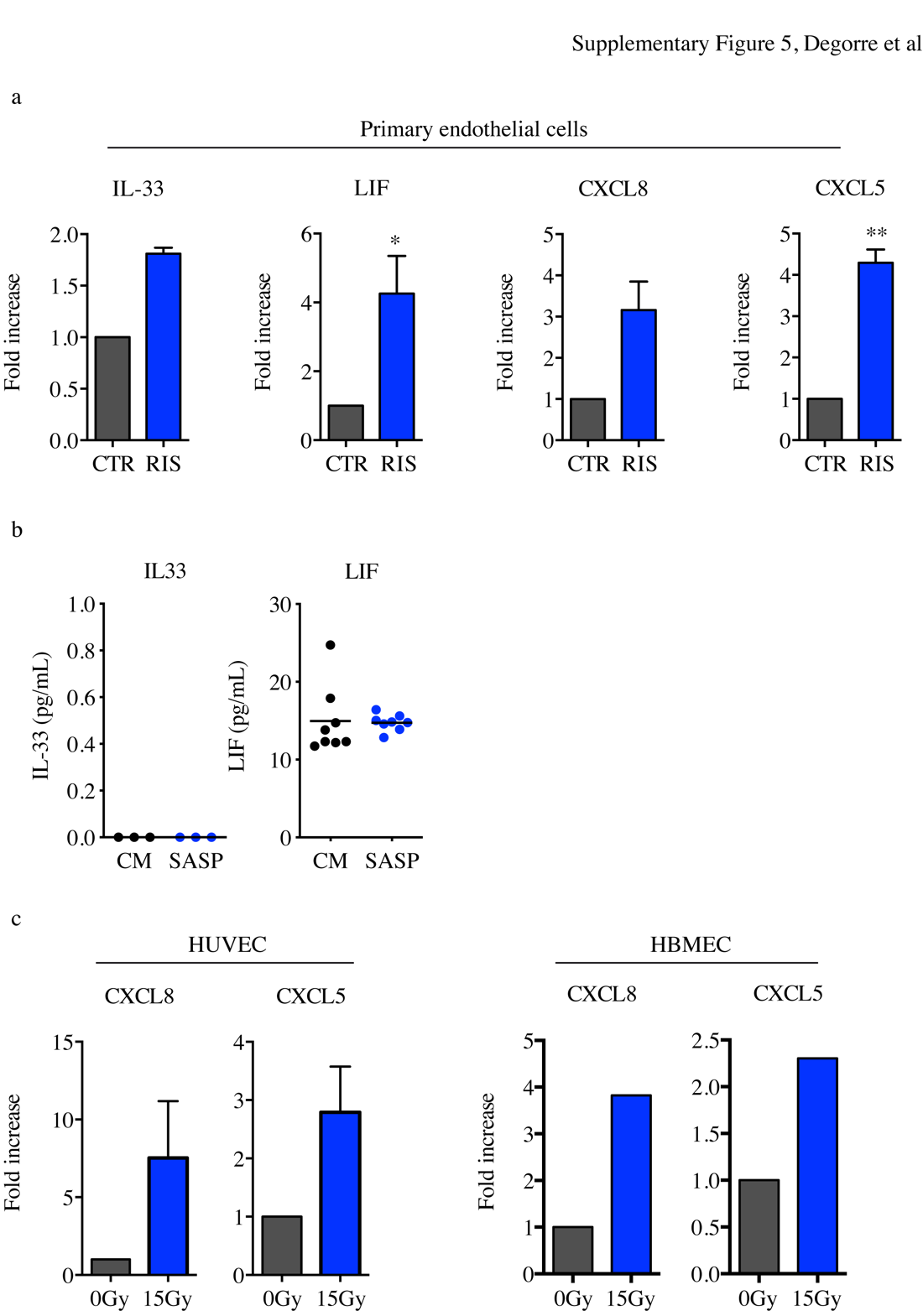

**Supplementary Figure 6 Establishment of R15CM-CXCL5/8, R15SASP-Mab and R15SASP-SB radiation-surviving U251 cells**.

Secretomes from primary endothelial HMVEC-L cells were collected from either unirradiated (CM) or RIS endothelial cells (SASP). Then U251 cells were plated at low density, preconditioned in either CM supplemented with CXCL8 and CXCL5 cytokines, SASP supplemented with CXCR2 monoclonal antibody MAb331 or SASP supplemented with SB3322235 CXCR2 antagonist before being irradiated at 15Gy. Three weeks later, R15CM-CXCL8, R15SASP-MAb and R15SASP-SB were collected and injected orthotopically in NSG mice.

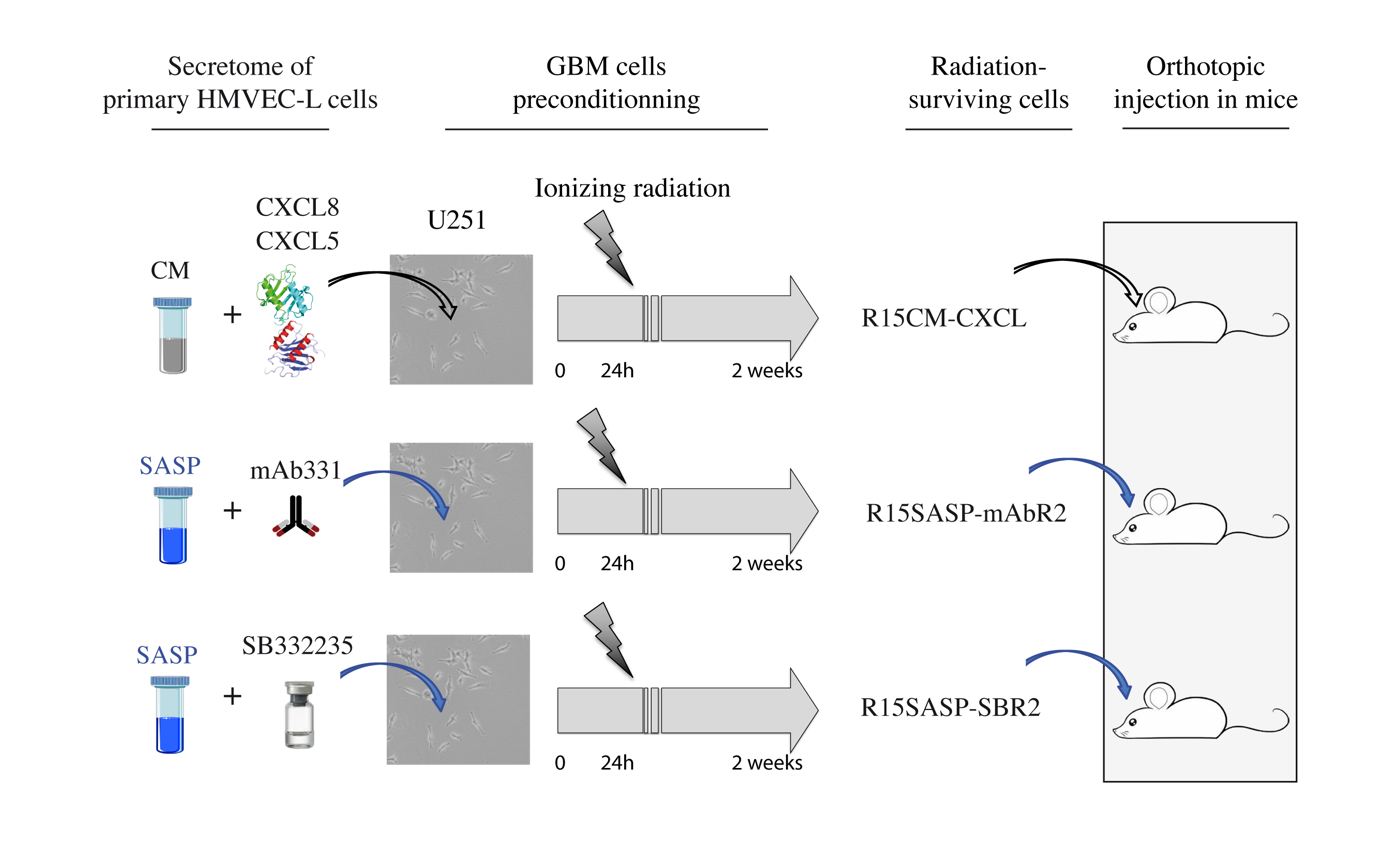
